# RTEL1 and MCM10 overcome topological stress during vertebrate replication termination

**DOI:** 10.1101/2022.01.27.478093

**Authors:** Lillian V. Campos, Briana H. Greer, Darren R. Heintzman, Tamar Kavlashvili, W. Hayes McDonald, Kristie Lindsey Rose, Brandt F. Eichman, James M. Dewar

## Abstract

Topological stress can cause replication forks to stall as they converge upon one another during termination of vertebrate DNA synthesis. However, replication forks ultimately overcome topological stress and complete DNA synthesis, suggesting that alternative mechanisms can overcome topological stress. We performed a proteomic analysis of converging replication forks that were stalled by topological stress in *Xenopus* egg extracts. We found that the helicase RTEL1 and the replisome protein MCM10 were highly enriched on DNA under these conditions. We show that RTEL1 normally plays a minor role during fork convergence while the role of MCM10 is normally negligible. However, RTEL1 and MCM10 both become crucially important for fork convergence under conditions of topological stress. RTEL1 and MCM10 exert non-additive effects on fork convergence and physically interact, suggesting that they function together. Furthermore, RTEL1 and MCM10 do not impact topoisomerase activity but do promote fork progression through a replication barrier. Thus, RTEL1 and MCM10 appear to play a general role in promoting progression of stalled forks, including when forks stall during termination. Overall, our data identify an alternate mechanism of termination involving RTEL1 and MCM10 that can be used to complete DNA synthesis under conditions of topological stress.

## INTRODUCTION

Eukaryotic DNA replication is organized into discrete steps: initiation, elongation, and termination (O’Donnell et al. 2013; Siddiqui et al. 2013; Bell and Labib 2016; Bleichert et al. 2017). This scheme occurs ∼60,000 times per S phase in human cells (Huberman and Riggs 1968) and must be executed faithfully to ensure genome stability. An excess of pre-replication complexes (pre-RCs) are loaded onto DNA but only a subset of these are activated (McIntosh and Blow 2012). This allows for defects in initiation or elongation to be overcome by activation of a proximal pre-RC, resulting in a new initiation event (Limas and Cook 2019). Additionally, elongation defects can be overcome by a wide array of DNA damage responses can aid fork progression (Cortez 2019). In contrast to initiation and elongation, much less is known about how obstacles to termination of DNA synthesis are overcome. This is important to address because it is well established that terminating replication forks are susceptible to stalling during SV40 replication (Tapper and DePamphilis 1978; Seidman and Salzman 1979) and recent data suggests the same is also true in eukaryotes (Devbhandari et al. 2017; Deegan et al. 2019).

Resolution of topological stress is particularly important during termination of DNA replication (Supplemental Fig. S1A) compared to earlier stages of replication. Type II topoisomerases are important to prevent converging replication forks from stalling during viral, bacterial, eukaryotic, and vertebrate replication termination (Ishimi et al. 1992; Hiasa and Marians 1996; Deegan et al. 2019; Heintzman et al. 2019). The unique role of type II topoisomerases is thought to reflect their ability to resolve intertwines of double-stranded DNA that form behind replication forks, (‘pre-catenanes’), which cannot be removed by other topoisomerases (Ullsperger et al. 1995; Vos et al. 2011; Pommier et al. 2016). Without type II topoisomerase activity, pre-catenanes accumulate either during termination (Champoux and Been 1980; Le et al. 2019) or throughout replication (Heintzman et al. 2019) and cause replication forks to stall due to topological stress (Supplemental Fig. S1B). However, replication forks can ultimately complete DNA synthesis without type II topoisomerases (Supplemental Fig. S1B) (Ishimi et al. 1992; Hiasa and Marians 1996; Baxter and Diffley 2008; Deegan et al. 2019; Heintzman et al. 2019). These data suggest that additional mechanisms exist that can overcome topological stress as replication forks converge.

Two mechanisms have been identified that promote resolution of converging forks independent of type II topoisomerases. In budding yeast, the 5’-3’ helicases Rrm3 and Pif1 both promote fork merger during replication termination (Deegan et al. 2019). These proteins, as well as their fission yeast paralog Pfh1, are also implicated in replication through, and termination at, replication barriers (Ivessa et al. 2000; Sabouri et al. 2012; Steinacher et al. 2012), suggesting that the obstacles posed by termination and replication barriers may be resolved by similar mechanisms. In bacteria, the RecQ helicase can cooperate with topoisomerase III to resolve converging forks independent of type II topoisomerase activity (Suski and Marians 2008). However, budding yeast Top3 is unable to promote resolution of converging forks (Deegan et al. 2019). Thus, type II topoisomerases and 5’-3’ helicases are the only proteins implicated in fork convergence in eukaryotes. However, it is unclear whether this is true in vertebrates and whether other proteins might promote fork convergence.

To identify proteins that might promote fork merger independently of type II topoisomerases, we performed a mass spectrometry analysis of proteins bound to DNA in TOP2α-immunodepleted *Xenopus* egg extracts. We found that the 5’-3’ helicase RTEL1 and the replisome component MCM10 were highly enriched on DNA under these conditions. We found that RTEL1 and MCM10 are normally dispensable for termination but become crucially important when forks stall during termination due to the absence of TOP2α. RTEL1 and MCM10 function together and co-IP, indicating that they form a single functional unit. RTEL1 and MCM10 do not affect utilization of TOP2α during DNA replication, indicating that they do not regulate pre-catenane generation, but are both important for fork progression through a replication barrier, independent of any effects on termination. Our data show that RTEL1-MCM10 facilitate fork progression through multiple obstacles to DNA replication and promote an alternate pathway for replication termination that allows replication forks to overcome topological stress.

## RESULTS

### A proteomic screen for proteins bound to terminating replication forks

To identify proteins that might promote fork merger independently of type II topoisomerases, we performed a proteomic analysis of proteins bound to replicating DNA when topoisomerase II (TOP2) was removed or inhibited. Under these conditions, replication forks stall during termination due to accumulation of topological stress (Heintzman et al. 2019). Plasmid DNA was replicated in mock- or TOP2α-depleted *Xenopus* egg extracts as previously described (Heintzman et al. 2019). In parallel, replication was performed in the presence of the TOP2 inhibitor ICRF-193 (‘TOP2-i’) as an alternate means of preventing TOP2 activity (Heintzman et al. 2019). Chromatinized plasmid DNA was recovered 18 minutes after the onset of DNA synthesis, when most forks have normally merged but are stalled when TOP2 activity is prevented (Fig. 1A) (Heintzman et al. 2019). Chromatin-bound proteins were then analyzed by chromatin mass spectrometry and quantified by label free quantification, as previously described (Räschle et al. 2015; Dewar et al. 2017; Larsen et al. 2019). This approach identified a total of 495 proteins (Supplemental Table S1). Importantly, TOP2α was ∼200-fold reduced in TOP2α-depleted conditions compared to the mock (Supplemental Fig. S1C), consistent with effective removal of TOP2α from the extracts. Additionally, TOP2α was ∼60% enriched in the presence of TOP2-i compared to the mock (Supplemental Fig. S1C), consistent with the ability of TOP2-i to trap TOP2 on DNA (Roca et al. 1994). Thus, this dataset contains proteins that are chromatin-bound during normal DNA replication and when TOP2 is either absent or inhibited.

**Figure 1:**
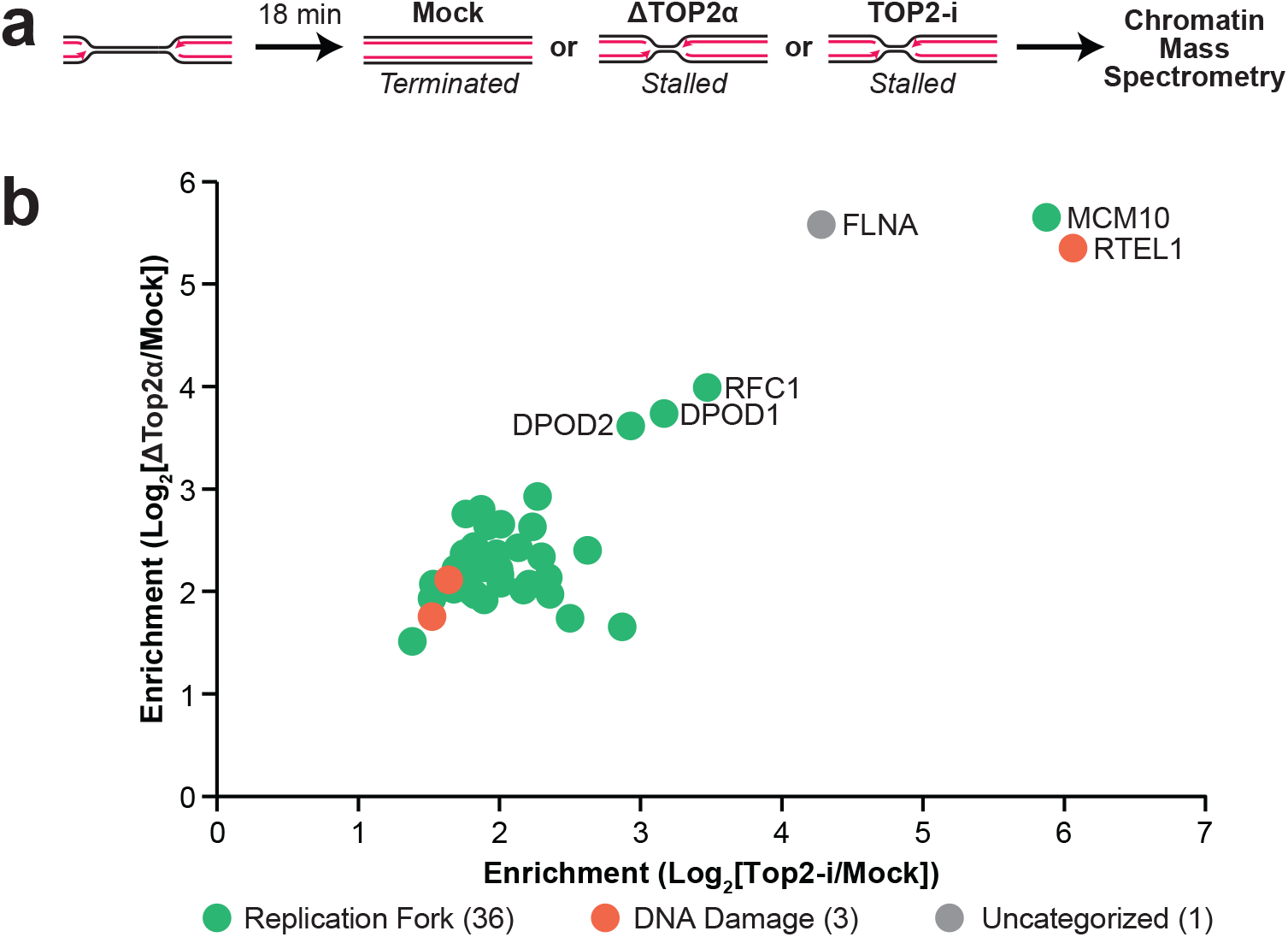
Proteomic analysis of replication forks stalled by topological stress. (A) Plasmid DNA was replicated in *Xenopus* egg extracts that were either mock-depleted (Mock), TOP2α-depleted (ΔTOP2α), or treated with TOP2 inhibitor ICRF-193 (TOP2-i). At 18 minutes, reactions were stopped and DNA-bound proteins were analyzed by Chromatin Mass Spectrometry and Label Free Quantification. (B) Proteins from (A) that were ≥2-fold enriched in ΔTOP2α and TOP2-i conditions compared to mock are plotted according to their enrichment. Each protein was manually assigned a category. The 6 most highly enriched proteins are labeled. Values were calculated from 3 independent experiments for each condition. See also Supplemental Fig. S1C and Supplemental Tables S1 and S2.

We next searched our dataset for proteins that exhibited increased DNA binding when TOP2 was either absent or inhibited. We selected proteins that were ≥2-fold enriched in TOP2α-depleted and TOP2-i treated conditions compared to the mock condition (Supplemental Fig. S1C, Table S2). We identified 40 proteins and in all cases enrichment was statistically significant (P≤0.05) for at least one of the two comparisons (Fig. 1B, Supplemental Fig. S1C).

We then categorized the proteins based on their previously reported functions. Most proteins (36) function at and/or travel with replication forks (Fig. 1B, Supplemental Fig. S1C), which is consistent with replication fork stalling during termination, which prevents removal of replication proteins from DNA (Heintzman et al. 2019). We also identified 3 proteins that are implicated in DNA damage responses (Fig. 1B, Supplemental Fig. S1C), which may reflect responses to replication fork stalling. We also identified Filamin-A (FLNA), which could not readily be categorized. FLNA promotes branching of actin filaments, which play a role during DNA replication in this extract system (Parisis et al. 2017). Additionally, FLNA has been implicated in the DNA damage response (Velkova et al. 2010). Thus, enrichment of FLNA could reflect a role in either DNA replication or DNA damage. Importantly, most replication fork and DNA damage proteins were ∼2-8 fold enriched in both conditions (Fig. 1B). However, 5 proteins were appreciably further enriched in both conditions. The PCNA loader RFC1 and the lagging strand DNA polymerase δ subunits DPOD1 and DPOD2 were ∼8-16 fold enriched. Enrichment of these proteins could reflect the switch from DNA synthesis by leading strand polymerase ε to polymerase δ, that was suggested to occur during replication termination (Zhou et al. 2019). The two remaining proteins, RTEL1 and MCM10, were ∼32-64 fold enriched. Thus, based on enrichment alone, RTEL1 and MCM10 were the strongest candidates revealed by our analysis.

RTEL1 is a 5’-3’ DNA helicase that can unwind DNA ahead of a stalled replication fork to help overcome obstacles to fork progression (Sparks et al. 2019). Additionally, analogous 5’-3’ helicases from yeast have been implicated in resolution of termination replication forks in cells and reconstituted systems, suggesting that the same may be true for RTEL1 (Ivessa et al. 2000; Sabouri et al. 2012; Steinacher et al. 2012; Deegan et al. 2019). MCM10 promotes DNA unwinding by the replicative helicase and is essential for replication initiation in yeast (Douglas et al. 2018). However, in vertebrates it plays an important but non-essential role in replication initiation (Wohlschlegel et al. 2002; Chadha et al. 2016). MCM10 is also implicated in replication fork stability in vertebrates, which could be important to overcome fork stalling during termination (Chadha et al. 2016). Thus, based on the reported activities of RTEL1 and MCM10 these proteins are ideally suited to promote resolution of terminating replication forks.

### RTEL1 and MCM10 promote replication fork merger

We next tested whether RTEL1 promotes replication termination. To this end we replicated plasmid DNA in mock-immunodepleted (‘mock-depleted’) extracts or RTEL1-immunodepleted (‘RTEL1-depleted’) extracts, then monitored the formation of the final products of replication as a read-out for termination. In mock-depleted extracts, the final products were circular monomers (Fig. 2A, CMs) as expected (Dewar et al. 2015). In RTEL1-depleted extracts formation of circular monomers was delayed (Fig. 2B, lanes 1-6 and 13-18) and there was no appreciable impact on total DNA synthesis (Supplemental Fig. S2A). Thus, the delay in circular monomer formation was due to a termination defect, rather than a defect during initiation or elongation of DNA synthesis. Importantly, θ structures (Fig. 2A, θs) but not catenanes (Fig. 2A, cats) persisted in RTEL1-depleted extracts (Fig. 2B, lanes 1-6 and 13-18) demonstrating that the delay in circular monomer formation was due to a fork merger defect. This allowed us to quantify circular monomer formation as a read-out for fork merger (Fig. 2C), which confirmed a modest delay in fork merger in RTEL1-depleted extracts. The RTEL1 antibody we used was previously verified to be specific to RTEL1 (Sparks et al. 2019), which demonstrates that the fork merger defect observed in RTEL1-depleted extracts is due to loss of RTEL1. Thus, RTEL1 promotes replication fork merger.

**Figure 2:**
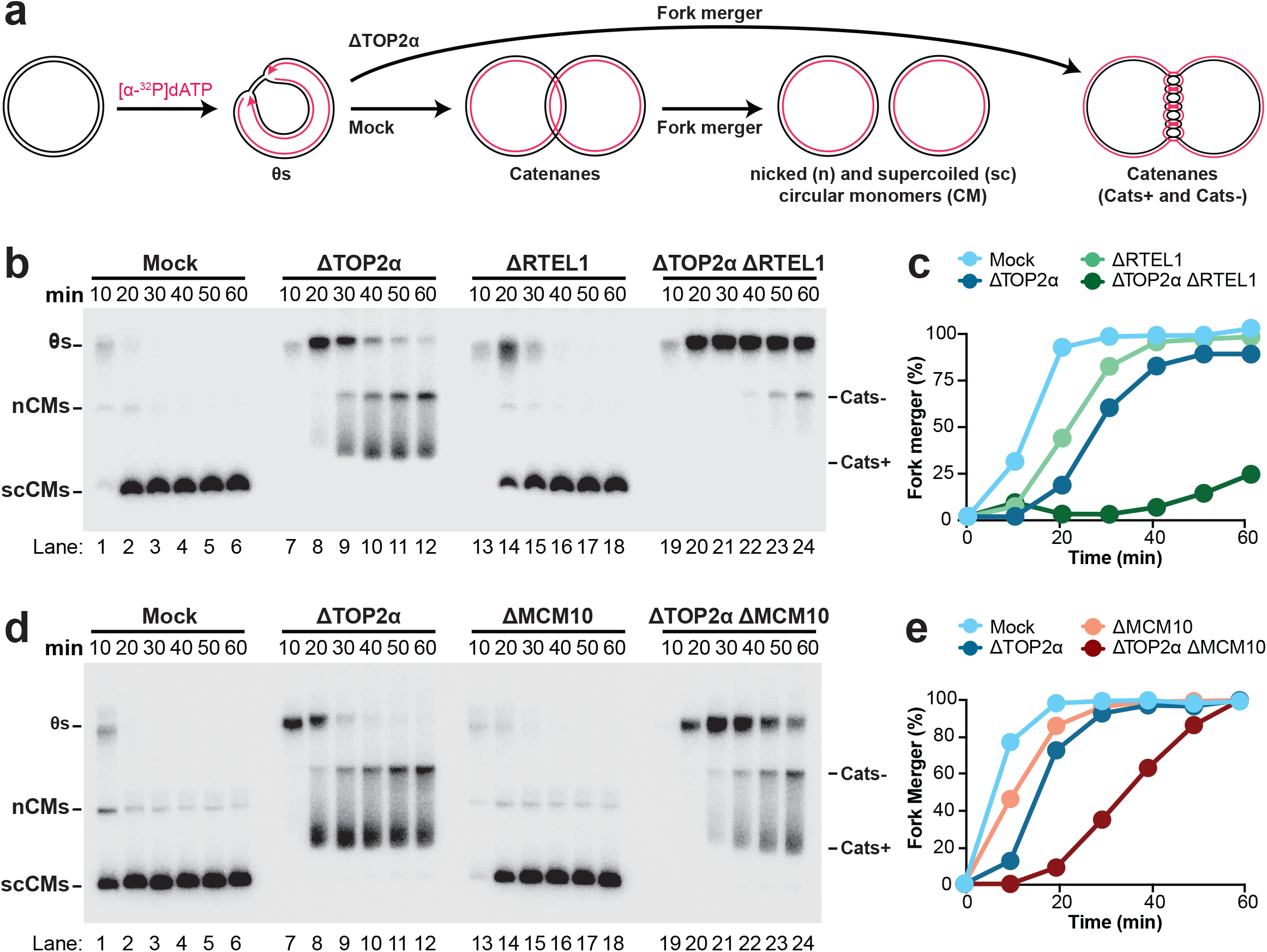
RTEL1 and MCM10 promote fork merger in the absence of TOP2α. (A) Plasmid DNA was replicated using Xenopus egg extracts in the presence of [α-^32^P]dATP to label newly-synthesized DNA strands under mock-depleted and ΔTOP2α-depleted conditions. The expected DNA structures, identified in (Heintzman et al. 2019), are indicated. (B) Replication was performed as in (A) with and without co-depletion of RTEL1. Samples were separated on a native agarose gel and visualized by autoradiography. (C) Quantification of fork merger from (B). See also Supplemental Fig. S2A. An independent experimental replicate is shown in Supplemental Fig. S2B. (D) Replication was performed as in (A) with and without co-depletion of MCM10. Samples were separated on a native agarose gel and visualized by autoradiography. (E) Quantification of fork merger from (D). See also Supplemental Fig. S2D. An independent experimental replicate is shown in Supplemental Fig. S2E.

RTEL1 was enriched on DNA when replication forks were stalled due to loss of TOP2 activity (Fig. 1A). To test whether RTEL1 was particularly important for fork merger under these conditions, we monitored the final products of replication in TOP2α-depleted and TOP2α-RTEL1-depleted (i.e. double depleted of both TOP2α and RTEL1) extracts. In TOP2α-depleted extracts the final products of replication were highly catenated catenanes (Fig. 2A, Cats+) which arise from fork merger and are slowly converted to less-catenated species (Fig. 2B, Cats-) by residual TOP2 activity (Heintzman et al. 2019). In TOP2α-RTEL1-depleted extracts formation of catenanes (Cats+ and Cats-) was dramatically impaired (Fig. 2B, lanes 7-12 and 19-24). Additionally, θ structures persisted at late time points in TOP2α-RTEL1-depleted extracts compared to TOP2α-depleted extracts (Fig. 2B, lanes 7-12 and 19-24). This demonstrated that the impaired formation of catenanes was due to a fork merger defect. We quantified catenane formation as a read-out for fork merger (Fig. 2C), which showed that the difference in fork merger between TOP2α-depleted and TOP2α-RTEL1-depleted extracts was much greater than between mock-depleted and RTEL1-depleted extracts. Thus, loss of RTEL1 led to a much greater fork merger delay in TOP2α-depleted extracts than in mock-depleted. Additionally, the fork merger delay in RTEL1-depleted extracts was less than in TOP2α-depleted extracts (Fig. 2C). These data show that RTEL1 ordinarily plays a more minor role than TOP2α during fork merger but becomes crucial when TOP2α activity is lost.

To address whether MCM10 also promotes fork merger we performed the same analysis as with RTEL1 (above). In MCM10-depleted extracts formation of circular monomers was modestly delayed (Fig. 2D, lanes 1-6 and 13-18) but this could be attributed to slowed replication (Supplemental Fig. S2D). Importantly, this delay had little impact on the fraction of circular monomers produced (Fig. 2E), indicating that there was negligible effect on termination. Instead, the delay in formation of circular monomers was due to delayed initiation of DNA replication, as previously described (Wohlschlegel et al. 2002; Chadha et al. 2016). In TOP2α-MCM10-depleted extracts formation of catenanes was dramatically impaired (Fig. 2D, lanes 7-12 and 19-24, Fig. 2E) and θ structures persisted at late time points (Fig. 2D, lanes 7-12 and 19-24), suggesting a fork merger defect. When the delay in replication in MCM10-depleted extracts was accounted for, a clear delay in fork merger was observed (Supplemental Fig. S2G), demonstrating that fork merger was delayed in TOP2α-MCM10-depleted extracts compared to TOP2α-depleted extracts. Importantly, the fork merger defect observed in TOP2α-MCM10-depleted extracts was rescued by re-addition of recombinant *Xenopus* MCM10 (Supplemental Fig. S2H-I), which confirmed that the fork merger defect was due to loss of MCM10. These data show that MCM10 normally plays a negligible role in promoting replication fork merger but becomes important when TOP2α activity is lost. Overall, our results show that RTEL1 and MCM10 are crucial to overcome topological stress during replication termination.

### RTEL1 and MCM10 function together

RTEL1 and MCM10 exhibited similar levels of binding in our proteomic analysis (Fig. 1) and were both crucial for fork merger following inactivation of Top2α (Fig. 2). To address whether RTEL1 and MCM10 function together we tested whether immunodepletion of both proteins together resulted in a non-additive effect compared to immunodepletion of either protein alone. This could not be done in TOP2α-depleted extracts because immunodepletion of three proteins would compromise extract function. Instead, we immunodepleted either or both proteins in the presence of a low dose of TOP2-i, which also inhibits fork merger (Heintzman et al. 2019). Low dose TOP2-i led to formation of several catenated species that migrated at a similarly to θ structures (Supplemental Fig. S3A, lanes 1-6), which made it challenging to measure products of fork merger. We therefore purified DNA intermediates and performed restriction digests to eliminate topoisomers (Fig. 3A). This allowed us to directly monitor fork merger by measuring conversion of double Y structures to the linear products of fork merger (Fig. 3A). Under these conditions, fork merger was largely complete by 60 minutes (Fig. 3B, lanes 1-6, 3C). RTEL1 immunodepletion delayed fork merger by ∼80 minutes (Fig. 3B, lanes 7-12, 3C) while MCM10 immunodepletion delayed fork merger by ∼20 minutes (Fig. 3B, lanes 13-18, 3C), consistent with a fork merger being more dependent on RTEL1 than MCM10 (Fig. 2). However, fork merger was not delayed following RTEL1-MCM10 immunodepletion compared to RTEL1 alone (Fig. 3B, lanes 7-12 and 19-24, 3C). Similar results were obtained when loss of θ structures was measured as a read-out for fork merger (Supplemental Fig. S3B). There was little impact on total DNA synthesis in all conditions tested (Supplemental Fig. S3C), indicating that these differences were due to defects in termination rather than earlier stages of DNA synthesis. Overall, these data show that MCM10 and RTEL1 function together to promote fork merger.

**Figure 3:**
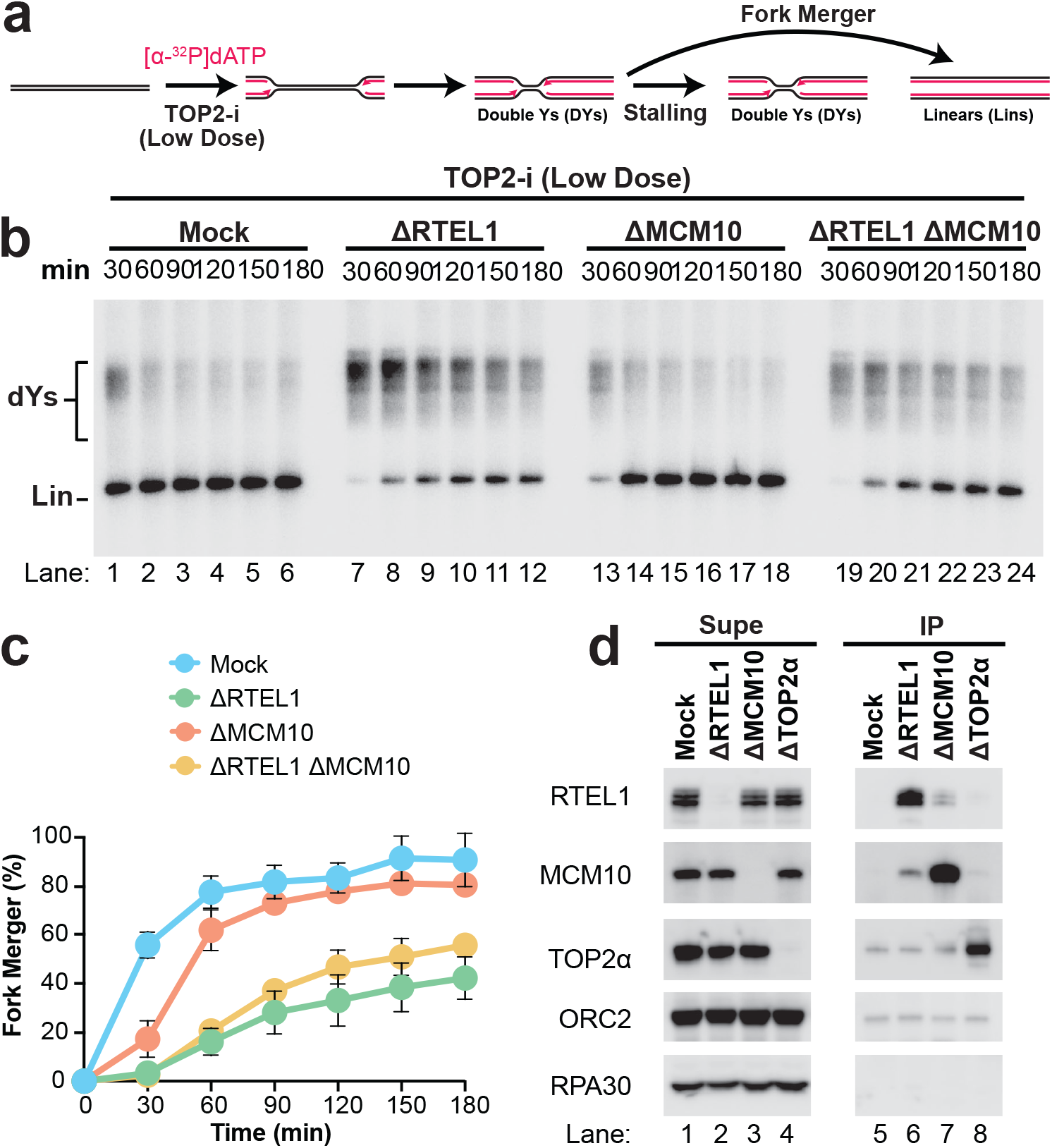
RTEL1 and MCM10 cooperate to promote fork merger. (A) Plasmid DNA was replicated in the presence of TOP2-i using extracts that were mock treated (mock) or depleted of RTEL1 (ΔRTEL1), MCM10 (ΔMCM10) or both (ΔRTEL1 ΔMCM10). Plasmids were purified and digested with XmnI, which cuts the plasmid once and allows unambiguous identification of replication fork structures (‘DYs’) and fully replicated molecules (‘Lins’) so that fork merger can be measured. (B) Replication was performed as in (A). Samples were separated on a native agarose gel and visualized by autoradiography. See also Supplemental Fig. S3A. (C) Quantification of fork merger from (B). Mean ± SD, n=3 independent experiments. See also Supplemental Fig. S3B-C. (D) *Xenopus* egg extracts were mock treated (mock) or depleted of RTEL1 (ΔRTEL1), MCM10 (ΔMCM10), or TOP2α (ΔTOP2α). The supernatant (Supe) and immunoprecipitate (IP) were then analyzed by Western blotting. The multiple RTEL1 bands correspond to multiple isoforms in Xenopus egg extracts, as previously reported (Sparks et al. 2019). A representative experiment of two independent repeats is shown.

Since MCM10 and RTEL1 function together we wondered whether they might also physically interact. To address this, we performed co-immunoprecipitation (co-IP) experiments and analyzed the immunoprecipitates by western blotting. Immunoprecipitation of RTEL1 led to co-IP of MCM10 (Fig. 3D, lanes 5 and 6) and *vice versa* (Fig. 3D, lanes 5 and 7). Co-IP was not detected between MCM10 or RTEL1 and ORC2 or RPA, which bind double- and single-stranded DNA, respectively (Fig. 3D, lanes 5-7). Thus, RTEL1 and MCM10 specifically interact, consistent with previous identification of MCM10 as an interacting partner of RTEL1 in murine cells (Vannier et al. 2013). Importantly, co-IP was not detected between either RTEL1 or MCM10 and TOP2α (Fig. 3D, lanes 5-7) and *vice versa* (Fig. 3D, lanes 5 and 8). Thus, RTEL1 and MCM10 physically interact, either directly or indirectly, but do not interact with TOP2α. These results show that at least two separate protein complexes promote replication fork merger (TOP2α and RTEL1-MCM10).

### MCM10 and RTEL1 do not limit the need for TOP2 activity

Immunodepletion of TOP2α causes replication forks to stall due to accumulation of catenanes (Heintzman et al. 2019). Thus, the role of RTEL1 and MCM10 in promoting fork merger could be explained by negative regulation of catenane formation, which would limit the need for TOP2α activity. We were unable to unambiguously measure catenanes as this can only performed on fully replicated molecules generated in the absence of TOP2 (Heintzman et al. 2019), but co-immunodepletion of either MCM10 or RTEL1 would interfere with formation of these same fully replicated molecules (Fig. 2). As an alternative strategy, we tested whether RTEL1 or MCM10 affect the abundance of TOP2:Cleavage Complexes (TOP2:CCs) generated by the TOP2 poison etoposide. TOP2:CCs arise when the TOP2 catalytic cycle is blocked and thus should act as a read out for TOP2 activity (Nitiss 2009). If RTEL1 or MCM10 normally negatively regulate catenane formation then loss of either protein would be expected to increase the abundance of TOP2:CCs.

We have previously shown that etoposide causes formation of TOP2:CCs during replication in Xenopus egg extracts (Van Ravenstein et al. 2021). To carefully quantify the formation of TOP2:CCs, we first replicated plasmid DNA in the presence of a low dose of etoposide (Fig. 4A). Reactions were stopped at different time points and mixed with a small amount of fully replicated control plasmid (pCTRL) that served as a loading control (Fig. 4A). Each reaction was then split in half and either left untreated or treated with proteinase K to remove TOP2:CCs (Fig. 4A). DNA was then subjected to phenol:chloroform extraction, which purified only DNA molecules that did not contain TOP2:CCs (Fig. 4A). The resultant DNA molecules were separated on native agarose gels to detect molecules that did not contain TOP2:CCs (Fig. 4B, – ProtK) or total DNA molecules (Supplemental Fig. S4A, +ProtK). This allowed us to calculate the percentage of molecules that contained TOP2:CCs (Fig. 4C). Etoposide treatment caused replication intermediates and nicked molecules to persist (Supplemental Fig. S4A, lanes 1-4,9-12). However, these molecules were lost when proteinase K treatment was omitted (Fig. 4B, lanes1-4,9-12), indicating that they contained TOP2:CCs. When we calculated the percentage of DPC-containing molecules we found that ∼75% of molecules contained DPCs and this declined by ∼2-fold over the course of our experiments. Thus, our experimental approach allowed us to monitor formation of a sub-saturating quantity of TOP2:CCs that were gradually resolved.

**Figure 4:**
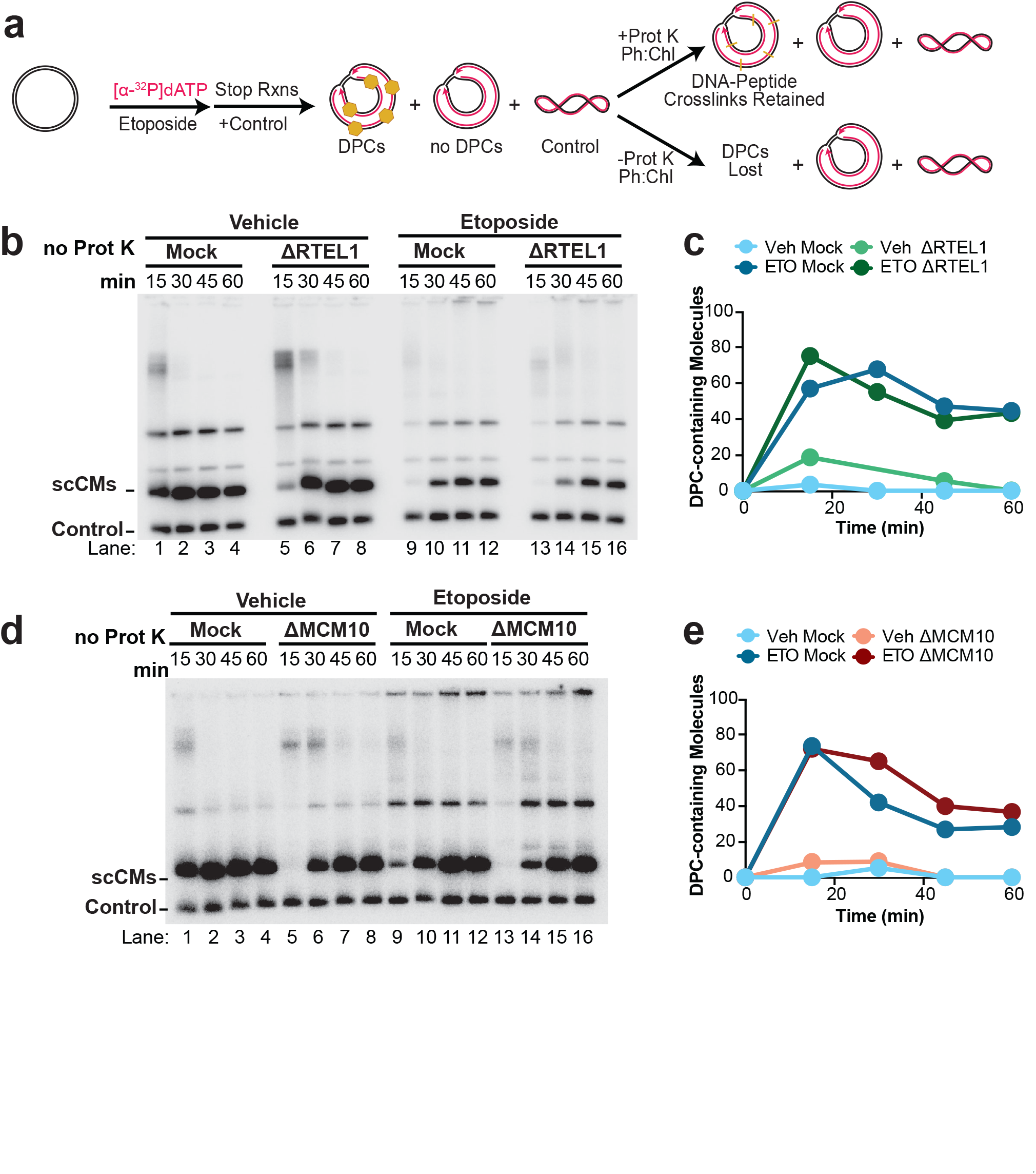
RTEL1 and MCM10 do not impact TOP2 utilization. (A) Plasmid DNA was replicated in the presence or absence of etoposide (ETO). Samples were combined with a radiolabeled control plasmid then split and treated with or without Proteinase K (Prot K). DNA lacking TOP2:CCs was then purified by phenol:chloroform extraction. (B) Replication was performed as in (A) with and without co-depletion of RTEL1. Samples that did not receive ProtK treatment were separated on a native agarose gel and visualized by autoradiography. ProtK treated samples are shown in Supplemental Fig. S4A. (C) Quantification of DPC-containing molecules from (B) and Supplemental Fig. S4A. See also Supplemental Fig. S4B. An independent experimental replicate is shown in Supplemental Fig. S4C. (D) Replication was performed as in (A) with and without co-depletion of MCM10. Samples that did not receive ProtK treatment were separated on a native agarose gel and visualized by autoradiography. ProtK treated samples are shown in Supplemental Fig. S4D. (E) Quantification of DPC-containing molecules from (D) and Supplemental Fig. S4D. See also Supplemental Fig. S4E. An independent experimental replicate is shown in Supplemental Fig. S4F.

To test whether RTEL1 or MCM10 affected the abundance of TOP2:CCs, we monitored their abundance in extracts that were either RTEL1-depleted (Fig. 4B-C, Supplemental Fig. S4A-C) or MCM10-depleted (Fig. 4D-E, Supplemental Fig. S4D-F). When proteinase K was omitted, the abundance of TOP2:CC-free DNA molecules was not appreciably reduced by either RTEL1 depletion (Fig. 4B, lanes 9-16) or MCM10 depletion (Fig. 4D, lanes 9-16) indicating that neither depletion appreciably increased abundance of TOP2:CCs. Quantification of TOP2:CC-containing molecules also revealed little difference between mock and depleted conditions following etoposide treatment (Fig. 4C,E). Although resolution of TOP2:CCs was delayed by ∼10 minutes following MCM10 depletion (Fig. 4E) this was likely due to the ∼10 minute delay in total DNA replication under these conditions (Supplemental Fig. S4E). Overall these data show that RTEL1 and MCM10 do not appreciably impact formation of TOP2:CCs following etoposide treatment and thus do not negatively regulate catenane generation.

### RTEL1 and MCM10 promote fork progression through a replication barrier

The requirement for RTEL1 and MCM10 following inactivation of TOP2α (Fig. 2) could reflect a general role for RTEL1 and MCM10 in responding to fork stalling. Accordingly, RTEL1 is important for fork progression when two forks converge upon a LacR-bound *lacO* array, which functions as a replication barrier (Sparks et al. 2019). However, it is unclear whether this reflects a general role in fork progression or a specific role during termination. Thus, we investigated the roles of RTEL1 and MCM10 in fork progression through a replication barrier, using a strategy that could determine whether any observed differences were due to fork progression or termination. We monitored the ability of replication forks to complete DNA synthesis within LacR arrays, which act as barriers to fork progression (Dewar et al. 2015). We compared fork merger in LacR-bound plasmids containing 16x (p[*lacO*x16]) or 32x (p[*lacO*x32]) copies of *lacO* (Fig. 5A). In both cases only a single termination event takes place. However, replication forks must overcome approximately twice as many bound LacR molecules in p[*lacO*x32] compared to p[*lacO*x16]. Thus, if RTEL1- or MCM10-depletion confers a termination defect then the delay compared to the mock condition should be similar for p[*lacO*x16] and p[*lacO*x32]. However, if a general fork progression defect is observed then the delay should be approximately twice as great in the depletion condition compared to the mock.

**Figure 5:**
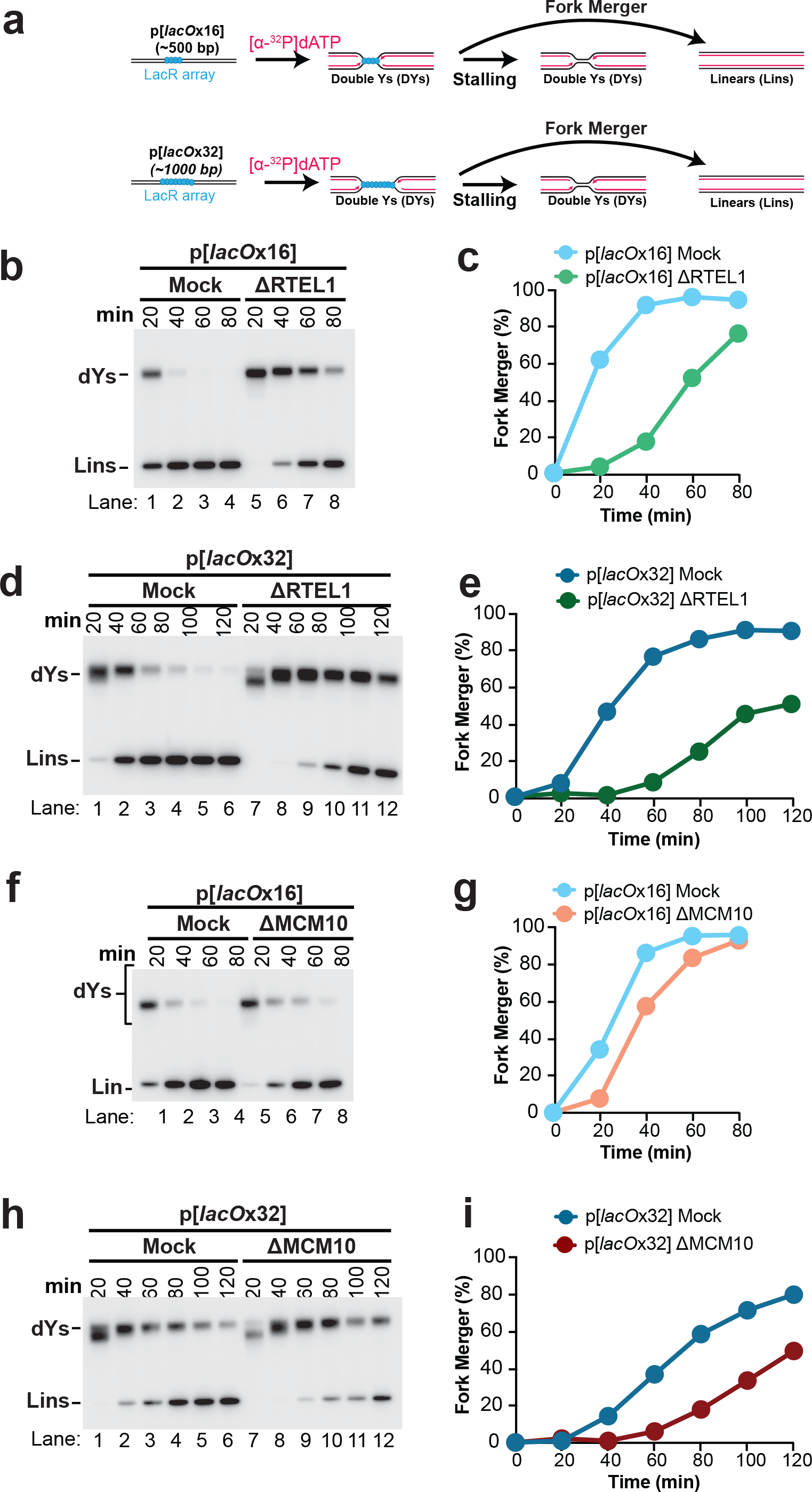
RTEL1 and MCM10 promote fork progression through a replication barrier. (A) Plasmid DNA harboring a 16x*lacO* array (p[*lacO*x16]) or a 32x*lacO* (p[*lacO*x32]) array was incubated with LacR to form a ∼250 bp or ∼500 bp replication barrier, respectively, then replicated. Purified replication intermediates were digested with XmnI, which cuts the plasmid once and allows unambiguous identification of replication fork structures (‘DYs’) and fully replicated molecules (‘Lins’) so that fork merger can be measured. (B) p[*lacO*x16] was replicated as indicated in (A) using mock-treated (Mock) and RTEL1-depleted (ΔRTEL1) extracts. Samples were separated on a native agarose gel and visualized by autoradiography. (C) Quantification of fork merger from (B). An independent experimental replicate is shown in Supplemental Fig. S5B. (D) As part of the experiment depicted in (B) p[*lacO*x32] was replicated using mock-treated (Mock) and RTEL1-depleted (ΔRTEL1) extracts.g (E) Quantification of fork merger from (D). An independent experimental replicate is shown in Supplemental Fig. S5C. (F) p[*lacO*x16] was replicated as indicated in (A) using mock-treated (Mock) and MCM10-depleted (ΔMCM10) extracts. Samples were separated on a native agarose gel and visualized by autoradiography. (G) Quantification of fork merger from (F). An independent experimental replicate is shown in Supplemental Fig. S5E. (H) As part of the experiment depicted in (F) p[*lacO*x32] was replicated using mock-treated (Mock) and RTEL1-depleted (ΔRTEL1) extracts. (I) Quantification of fork merger from (H). An independent experimental replicate is shown in Supplemental Fig. S5F.

We first tested the role of RTEL1 in promoting fork progression through a LacR array. Replication of p[*lacO*x16] and p[*lacO*x32] led to formation of several catenated species (Fig. Supplemental Fig. S5A) so restriction digests were performed on purified DNA to allow unambiguous quantification of fork merger (Fig. 5A, as in Fig. 3A). Double-Ys to persisted during replication of both p[*lacO*x16] (Fig. 5B, lanes 5-8, 5C) and p[*lacO*x32] (Fig. 5D, lanes 5-8, 5E) in RTEL1-depleted extracts, as expected (Sparks et al. 2019). Importantly RTEL1-depletion caused a ∼30 minute delay in fork merger during replication of p[*lacO*x16] and a ∼60 minute delay during replication of p[*lacO*x32] (Supplemental Fig. S5D). Because the extent of the delay is approximately doubled when the size of the LacR array is doubled this shows that RTEL1 plays a general role in promoting fork progression through a replication barrier that is not confined to termination.

We next tested the role of MCM10, as described for RTEL1 (above). Fork merger was also delayed during replication of both p[*lacO*x16] (Fig. 5F, lanes 5-8, 5G) and p[*lacO*x32] (Fig. 5H, lanes 5-8, 5I) in MCM10-depleted extracts. MCM10-depletion caused a ∼10 minute delay in fork merger during replication of p[*lacO*x16] and a ∼40 minute delay during replication of p[*lacO*x32] (Supplemental Fig. S5G). Thus, MCM10 also plays a general role in promoting fork progression through a replication barrier that is not confined to termination. Because MCM10 depletion delayed fork merger ∼4-fold during replication of p[*lacO*x16] compared to p[*lacO*x32] this also suggested that MCM10 became more important as the duration of the stall increased. Importantly, none of these effects could be attributed to slowed replication in MCM10 depleted extracts because the ∼10 minute delay in replication (Supplemental Fig. S2D,F) had negligible impact on these experiments, where reactions were sampled every 20 minutes (Supplemental Fig. S5H-J). Overall, these data show that MCM10 plays a general role in promoting fork progression through a replication barrier that is not confined to termination. Thus, both RTEL1 and MCM10 promote fork progression through replication barriers, independent of any role in termination.

## DISCUSSION

Our data show that RTEL1 and MCM10 are important for fork merger during termination of vertebrate DNA synthesis. These data support a new model for vertebrate replication termination (Fig. 6). Under normal circumstances efficient resolution of catenanes by TOP2 ensures that replication forks readily merge (Fig. 6Ci) irrespective of RTEL1 and MCM10 activity. If TOP2 is unable to fulfill this role then replication forks stall (Fig 6Cii) and fork merger occurs slowly, promoted by RTEL1 and MCM10, which work together to ensure continued progression of stalled forks and eventual fork merger. RTEL1 and MCM10 promote an alternative mechanism for replication termination that becomes crucially important when the normal TOP2-dependent mechanism fails. Thus, our data show that vertebrate termination can take place through multiple mechanisms and implicate a core replisome protein, MCM10, in this process.

**Figure 6:**
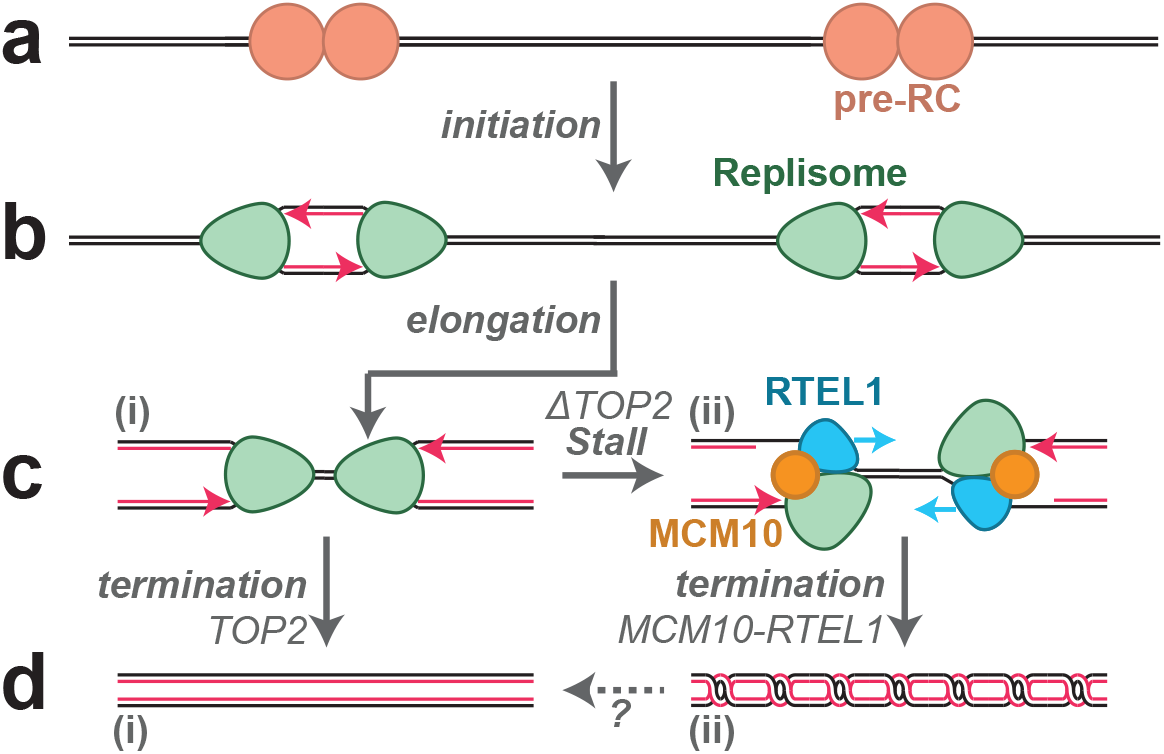
Model for vertebrate DNA replication. (A) Pre-RCs are loaded onto DNA prior to S phase (B) During S phase, pre-RCs are activated and associate with additional proteins to form replisomes that unwind DNA as they move away from each other in opposite directions (‘initiation’). Activated replisomes then establish replication forks that duplicate DNA until they encounter an opposing fork (‘elongation’). (C) (i) When two replication forks converge upon the same stretch of DNA (‘termination’) they complete DNA synthesis without stalling or slowing, provided that TOP2 activity prevents accumulation of topological stress. (ii) In the absence of TOP2 activity, converging replication forks stall due to accumulation of topological stress and RTEL1-MCM10 facilitate fork merger independently of TOP2. (D) (i) The final products of replication termination are normally fully replicated and unlinked. (ii) If forks stall due to lack of TOP2 activity (Fig. 6Cii) these products contain DNA catenanes. The fate of these structures and whether catenanes can be removed independently of TOP2 is unclear.

RTEL1-MCM10 promote fork merger in the absence of TOP2 activity. However, under these conditions the final products of replication are catenanes (Fig 6dii). These structures would need to be resolved to allow chromosome separation during mitosis. TOP2-independent mechanisms of unlinking catenanes have not been described, so it is likely that these species would undergo chromosome breakage upon exit from mitosis (Umbreit et al. 2020). It will be interesting to address whether this is the case or whether any additional TOP2-independent mechanisms exist to promote separation of catenated sister chromatids.

Maximal replication fork merger in the budding yeast reconstituted system requires both 5’-3’ helicase activity, supplied by either Pif1 or Rrm3, and topoisomerase activity, provided by both Top1 and Top2 (Deegan et al. 2019). Our data similarly implicate the 5’-3’ helicase RTEL1 and the topoisomerase TOP2 in fork merger during vertebrate replication termination. However, the importance of topoisomerases and 5’-3’ helicases differs between budding yeast and vertebrates. In the budding yeast reconstituted system and yeast cells Top2 makes a minor contribution to fork merger (Baxter and Diffley 2008; Deegan et al. 2019). Although another study reported a more significant role for Top2 during fork convergence in yeast cells (Fachinetti et al. 2010) this was only for a minority of termination events (McGuffee et al. 2013), consistent with a minor role for Top2. In contrast, loss of TOP2α causes essentially all forks to stall during vertebrate replication termination (Heintzman et al. 2019), indicating a major role for TOP2α. For 5’-3’ helicases priorities appear reversed and in budding yeast Rrm3 is normally crucial for termination (Deegan et al. 2019), while in vertebrates RTEL1 normally plays a minor role (Fig. 2). Thus, although similar mechanisms promote fork merger in both the yeast and vertebrate systems the priority placed on these mechanisms differs between systems. This may suggest that the underlying obstacles to fork merger differ between organisms.

Our experiments were performed using circular chromatinized minichromosomes that lack chromosomal features (e.g. centromeres) and many aspects of chromosome metabolism (e.g. transcription). It will therefore be important to test this model within the context of full-length chromosomes in cells. However, MCM10 is involved in replication initiation (Wohlschlegel et al. 2002; Chadha et al. 2016) and RTEL1 is required to replicate a variety of chromosomal features (Vannier et al. 2014). Thus, it will be difficult to address whether any under-replication of DNA is due to a termination defect rather than any earlier stage of DNA replication. We note that both RTEL1 and MCM10 promote fork progression during telomere replication (Vannier et al. 2012; Baxley et al. 2021), suggesting that both proteins might cooperate in other processes beyond termination. It will be interesting to determine which other cellular processes involve both RTEL1 and MCM10 and whether the two proteins cooperate.

To our knowledge, our study is the first to report functional cooperation between RTEL1 and MCM10. Because RTEL1 exerts a much stronger effect than MCM10 (Fig. 3C) our favored model is that RTEL1 fulfills some sort of role that is supported by MCM10. The most likely role for RTEL1 is unwinding DNA ahead of stalled replication forks, as previously described (Sparks et al. 2019). This is the simplest way that RTEL1 could contribute to fork progression given that no other enzymatic activities beyond its 5’-3’ helicase activity have been identified (Vannier et al. 2014). In contrast, it is unclear how MCM10 might assist RTEL1 given the multiple reported functions and protein-protein interactions of MCM10 (Baxley and Bielinsky 2017). Because RTEL1 and MCM10 interact (Fig. 3d) (Vannier et al. 2013), MCM10 may directly recruit RTEL1. However, most RTEL1 does not interact with MCM10 and *vice versa* so we disfavor this model. Alternatively, MCM10 may promote replisome stability, as previously reported (Chadha et al. 2016). This activity may allow MCM10 to help the replisome withstand forces experienced during fork stalling. Moreover, if RTEL1 were directly connected to the replisome then MCM10 might act as a ‘brace’, which could allow RTEL1 to exert more force in the direction of the replication obstacle. Yet another model is that MCM10 inhibits reannealing of the parental strands, as previously described (Wasserman et al. 2019). This is particularly appealing because topological stress would be expected to favor re-annealing of the parental DNA strands. It the future it will be important to understand how RTEL1 and MCM10 cooperate to promote fork progression.

## METHODS

### DNA replication in *Xenopus* egg extracts

*Xenopus* egg extracts were prepared from *Xenopus laevis* wild-type male and female frogs (Nasco) as previously described (Heintzman et al. 2019). Animal care protocols were approved by the Vanderbilt Division of Animal Care (DAC) and Institutional Animal Care and Use Committee (IACUC). Plasmid DNA (12.3 ng/μl final concentration) was incubated for 30 minutes with High Speed Supernantant (HSS) supplemented with nocodazole (3 ng/μl final concentration) and ATP regenerating system (ARS, final concentration: 20 mM phosphocreatine, 2 mM ATP and 5 ng/μl phosphokinase). Replication was initiated by addition of 2 volumes NucleoPlasmic Extract (NPE) supplemented with ARS, and DTT (2 mM final concentration). Reactions were stopped by addition of 10 volumes Stop solution (50 mM Tris-HCl pH 8, 25 mM EDTA, 0.5% SDS (w/v)) then treated with RNase (190 ng/ul final concentration) and Proteinase K (909 ng/ul).

To stall forks at a LacR array plasmid DNA was pre-incubated with LacR, as previously described (Dewar et al. 2015). ICRF-193 (TOP2-i) was dissolved in DMSO and added at a final concentration of 200 μM (Figure 1) or 25 μM (Figure 3). Etoposide (ETO) was dissolved in DMSO and added at a final concentration of 1.11 μM.

### Analysis of replication intermediates

To analyze replication intermediates, replication reactions were supplemented with [α-^32^P]dATP (final concentration: 67-240 nM) to radiolabel nascent DNA strands. In most cases, samples were directly analyzed by native agarose gel electrophoresis. To analyze fork merger, DNA was purified, then treated with XmnI (0.4 U/μl) for 1hour at 37°C, as previously described (Dewar et al. 2015), prior to gel electrophoresis. In all cases gel electrophoresis was performed using a native agarose gel (1% (w/v)) at 5V/cm in TBE (1X).

### Plasmid constructs

pJD156 (p[*lacO*x32]) was used for all experiments unless otherwise specified. pJD145 (CTRL, Figure 5), pJD152 (p[*lacO*x16]), pJD156 (p[*lacO*x32]) were previously described (Dewar et al. 2015).

### Protein purification

Biotinylated LacR protein was purified from *Escherichia coli* as previously described (Dewar et al. 2015). Full length *Xenopus* MCM10 was also purified from baculovirus-infected SF9 insect cells as previously described (Du et al. 2013) except with the following modifications; HiTrap Heparin HP (Cytivia) was used in place of Source Q for ion exchange; and the size exclusion step was omitted.

### Antibodies

Antibodies targeting *Xenopus* TOP2α, RTEL1, RPA, and ORC2 were previously described (Dewar et al. 2017; Heintzman et al. 2019; Sparks et al. 2019). The MCM10 antibody was generated in this study against a purified polypeptide of *Xenopus* MCM10 composed of the following sequence: SYSGHVPKKMARGANGLRERLCQGGFHYGGVSSMAYAATLGSTTAPKKTVQSTLSNMVVRGAEAIALEARQKIAAAKNVVQTDEFKELMTLPTPGALNLKKHLSGVSPQANCGKEGQPIQSISASTLLKQQKQQMLNARKKRAEESQKRFLESTEKSEKSSTLTSSACSVFQSPKQGAEFPNAQKMATPKLGRGFAEGDDVLFFDISPPPAPKLSTSAEAKKLLAIQKLQAKGQTLAKTDPNSIKRKRGSSS

### Immunodepletion and rescue

Immunodepletion of TOP2α, MCM10, and RTEL1 was performed as previously described for TOP2α (Heintzman et al. 2019). For co-depletion of two proteins, beads were combined for each round of depletion and the number of depletion rounds remained constant. In all experiments beads bound to control IgGs were used to ensure that the concentration beads and IgGs was identical for all conditions. For rescue experiments, xMcm10 was added to a final concentration of 100 ng/μl.

### Chromatin Mass Spectrometry

Chromatin mass spectrometry (‘CHROMASS’) (Räschle et al. 2015) was performed using modifications to analyze plasmid DNA (Larsen et al. 2019). Plasmid DNA was replicated in mock-treated extracts containing ICRF-193 (TOP2-i) or vehicle control (DMSO). In parallel plasmid DNA was replicated in TOP2α-depleted extracts containing vehicle control. At 18 minutes, plasmid DNA and associated proteins were captured on beads and then processed for CHROMASS and captured on C18 resin using STAGE tips (Nikkyo Technos). The resultant material was then analyzed by liquid chromatography-tandem mass spectrometry and quantified using label-free quantification, as outlined below. For each condition 3 experimental replicates were analyzed.

### Liquid chromatography-tandem mass spectrometry (LC-MS/MS) analysis

Peptides were eluted from STAGE tips with 70% acetonitrile, eluates were dried by speed vac centrifugation, and peptides were reconstituted in 6 μL of 0.1 % formic acid for analysis by LC-coupled tandem mass spectrometry (LC-MS/MS). An analytical column was packed with 20cm of C18 reverse phase material (Jupiter, 3 μm beads, 300Å, Phenomenox). Peptides (2.5 μL) were loaded on the reverse phase analytical column (360 μm O.D. x 100 μm I.D.) using a Dionex Ultimate 3000 nanoLC and autosampler. The mobile phase solvents consisted of 0.1% formic acid, 99.9% water (solvent A) and 0.1% formic acid, 99.9% acetonitrile (solvent B). Peptides were gradient-eluted at 350 nL/min using a 95-minute gradient. The gradient was as follows: 1-78 min, 2-40% B; 78-85 min, 40-95% B; 85-86 min, 95% B; 86-87 min, 95-2% B; 87-95 min (column re-equilibration), 2% B. Peptides were analyzed using a data-dependent method on a Q Exactive Plus (Thermo Scientific) mass spectrometer, equipped with a nanoelectrospray ionization source. The instrument method consisted of MS1 using an AGC target value of 3e6, followed by up to 20 MS/MS scans of the most abundant ions detected in the preceding MS scan. The MS2 AGC target was set to 5e4, dynamic exclusion was set to 20s, HCD collision energy was set to 27 nce, and peptide match and isotope exclusion were enabled. MS/MS spectra were searched against a *Xenopus laevis* database with peptide matches being integrated from MS1 scans, collated to the protein level, and then normalized using MaxQuant-LFQ v1.5.2.8 (Cox et al. 2014). All LFQ values are reported in Supplemental Table S1. Raw files and MaxQuant output tables are available at ProteomeXchange (http://proteomecentral.proteomexchange.org; ID = PXD PXD031170; Username; Password: 9RlqJLFR).

### Statistical analysis of Label Free Quantification

Resulting protein quantitative outputs were compared pairwise using ProStar (live.prostar-proteomics.org) (Wieczorek et al. 2017). All calculations were based on the LFQ intensities reported by MaxQuant. To identify proteins that were significantly up-regulated in TOP2-i treated or TOP2α-depleted conditions compared with control reactions a modified t-test was performed with Benjamini-Hochberg correction for multiple comparisons applied. All test results are reported in Supplemental Table S2.

### DNA-Protein Crosslink (DPCs) Detection Assay

DNA replication reactions were stopped by addition of 10 volumes Stop solution then treated with RNase, as outlined above. 0.2 volumes of radiolabeled control plasmid (pJD145, 0.25 ng/μl) was added to each sample, which was then split in two. Half the reaction was then treated with Proteinase K (889 ng/μl) while the other half was treated with vehicle control (de-ionized previously described (Dewar et al. 2015). Samples were then separated by gel electrophoresis and visualized by autoradiography to detect total DNA (Proteinase K treated samples) or non-crosslinked DNA (samples not treated with Proteinase K). For each sample, fraction of DNA molecules containing DPCs was calculated using the total lane signal for non-crosslinked DNA (ncDNA) and total DNA (totDNA) normalized to control plasmid (CTRL) as follows: [(totDNA/CTRL) – (ncDNA/CTRL)] / (totDNA/CTRL) x 100%

## ACKNOWLEDGEMENTS

J.M.D. was supported by NIH grant R35GM128696. B.F.E. was supported by NIH grant R35GM136401. Jesus Mallol Diaz provided experimental assistance.

---

**Supplemental Figure 1:**
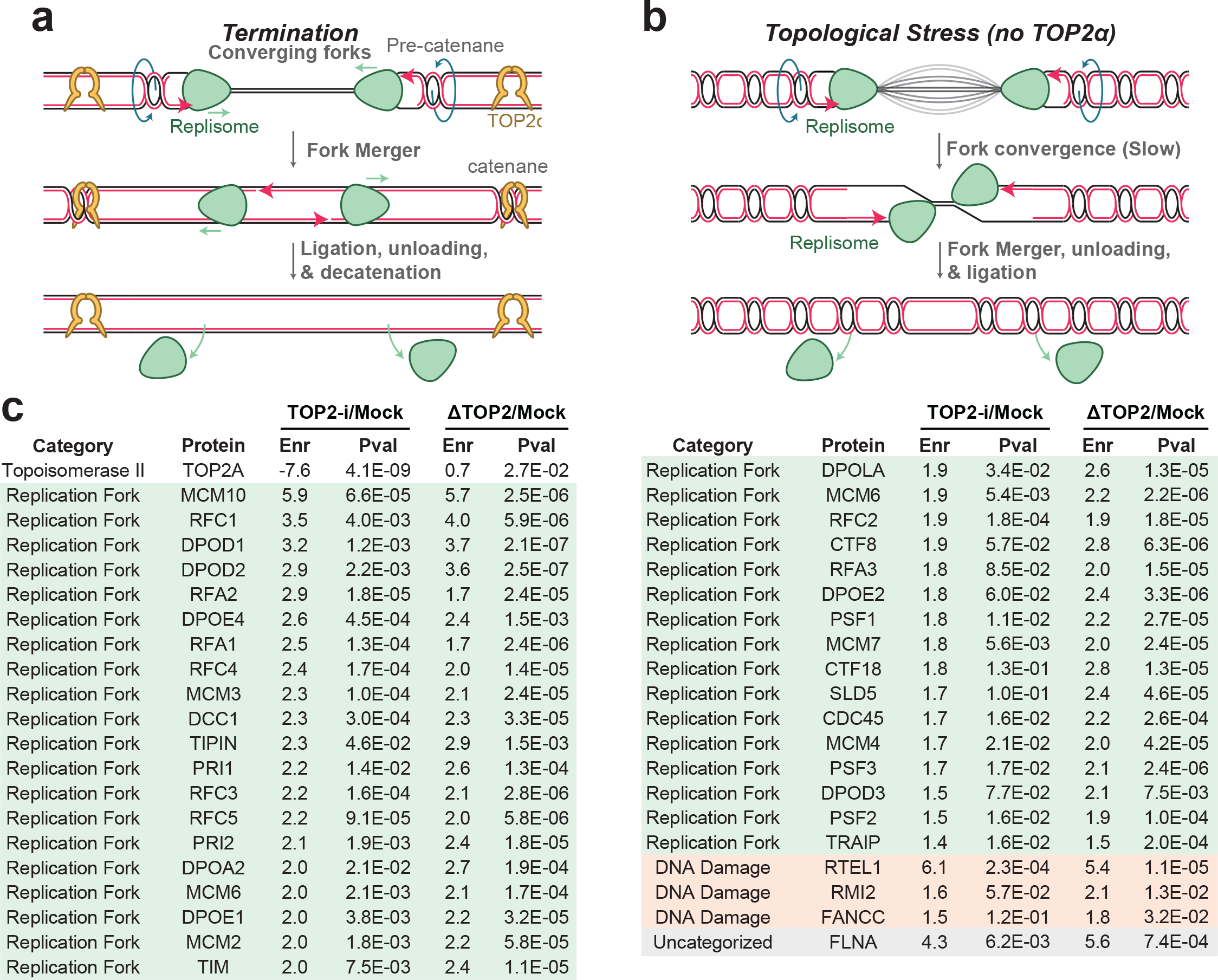
Effects of topological stress on replication termination. (A) Model for vertebrate replication under normal conditions (Dewar et al. 2015). Termination occurs when two converging replication forks encounter each other upon the same stretch of DNA (‘fork convergence’). Replisomes move past each other to unwind any remaining duplex (‘fork merger’) because the replicative helicases encircle opposite DNA strands. After forks merge, replisomes pass over fully replicated DNA from the opposing fork which allows any remaining daughter strand gaps to be rapidly filled in and ligated (‘ligation’). Replisomes are also removed by an active unloading pathway (‘unloading’) (Moreno et al. 2014; Dewar et al. 2015; Dewar et al. 2017; Sonneville et al. 2017). Pre-catenanes are formed throughout replication and resolved by Topoisomerase IIα (TOP2α). After fork merger, any pre-catenanes that were not resolved during replication form catenanes, which are resolved by TOP2α (‘decatenation’) (Heintzman et al. 2019). (B) In the absence of (TOP2α) converging replication forks stall during vertebrate replication termination, which delays all downstream termination events (Heintzman et al. 2019). Replication forks eventually unwind the remaining DNA duplex independent of TOP2α, which allows for fork merger, unloading, and ligation, and decatenation to take place. However, decatenation cannot take place due to the lack of TOP2 activity. (C) Full list of proteins from Figure 1B including their enrichment (Enr) and P-values (Pval) for different comparisons. Enr is a log2 transformation of the mean LFQ value from 3 independent experiments of ΔTOP2α or TOP2-i divided by the mean LFQ value from 3 independent experiments of mock. Pval was calculated using a modified t-test in ProStar performed with Benjamini-Hochberg correction for multiple comparisons. See also Supplemental tables S1 and S2.

**Supplemental Figure 2:**
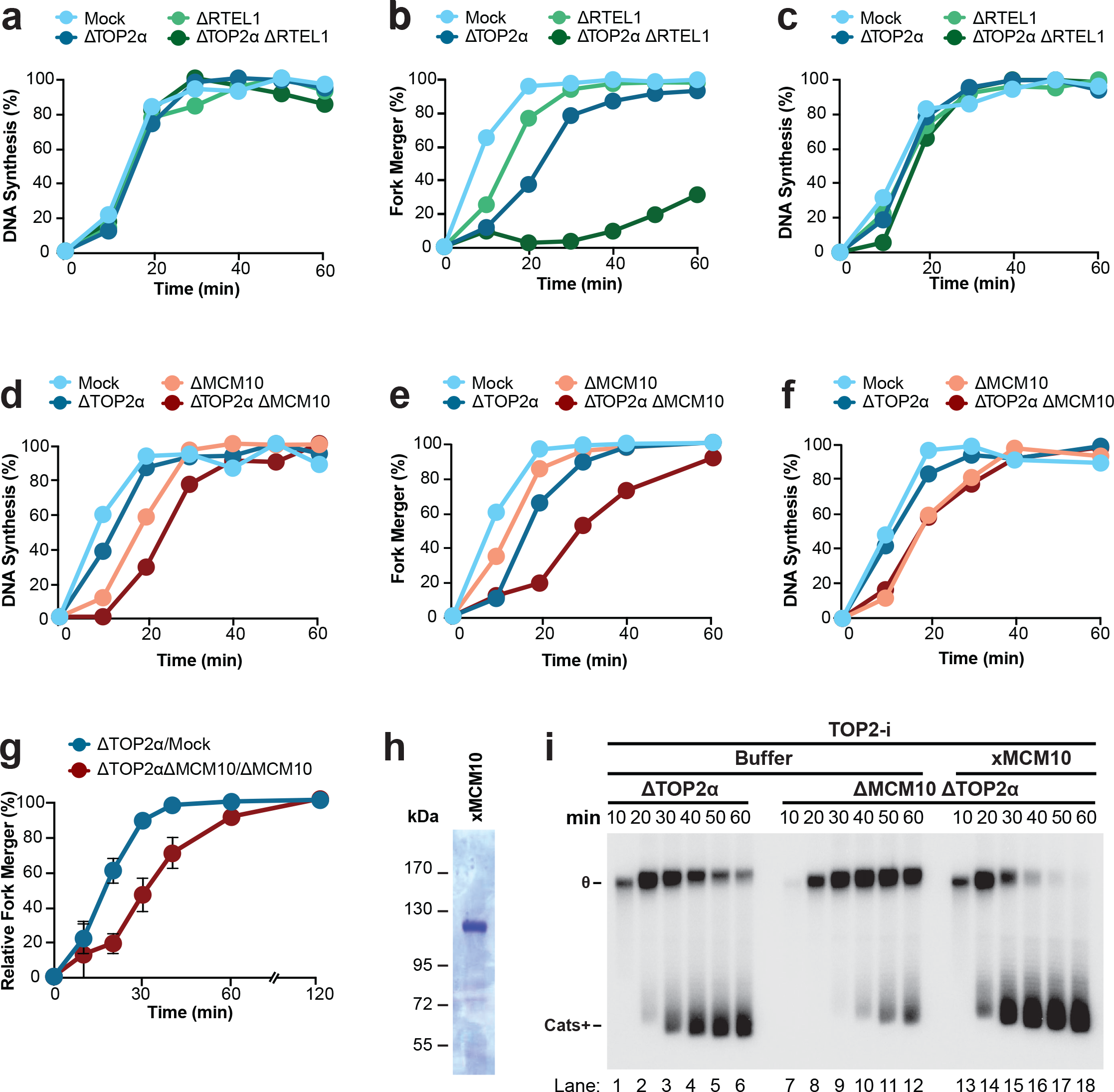
Contribution of RTEL1 and MCM10 to fork merger. (A) Quantification of total DNA synthesis from Figure 2B. (B) An experimental replicate of Figure 2C. (C) An experimental replicate of (A). (D) Quantification of total DNA synthesis from Figure 2D. (E) An experimental replicate of Figure 2E. (F) An experimental replicate of (D). (G) Slow replication in MCM10-depleted extracts could explain delayed fork merger in TOP2α-MCM10-depleted extracts compared TOP2α-depleted extracts in Figure 2E. To address this, fork merger in TOP2α-depleted extracts was calculated relative to mock conditions (ΔTOP2α/mock) and also in TOP2α-MCM10-depleted extracts relative to MCM10-depleted extracts (ΔTOP2α ΔMCM10/ΔMCM10) for 3 independent experiments. Relative fork merger for ΔTOP2α ΔMCM10/ΔMCM10 was delayed compared to ΔTOP2α/mock. Thus, MCM10 promotes fork merger in TOP2α-depleted extracts independent of replication slowing caused by MCM10-depletion. Mean ± SD, n=3 independent experiments. (H) SDS-PAGE gel of purified *Xenopus* MCM10. (I) To formally test whether the effects of MCM10 depletion in Figure 2D-E were caused by loss of MCM10, a rescue experiment was performed. Plasmid DNA was replicated as in Figure 2A in TOP2α-depleted and TOP2α-MCM10-depleted extracts. The latter was supplemented with either buffer control or purified MCM10 from (H). To simplify interpretation of the data, all extracts were supplemented with a low dose of Top2-i (25 μM) to block conversion of Cats+ to Cats-(Fig 2A-B). θ structures persisted in TOP2α-MCM10 depleted extracts compared to TOP2α depleted extracts (lanes 1-12) and this was rescued by addition of purified MCM10 (lanes 13-18). Thus, the fork merger defect conferred by MCM10 depletion was due to lack of MCM10 activity. MCM10 depletion also caused a delay in replication (compare lanes 1-2 and 7-8) and this was also rescued by addition of purified MCM10 (compare lanes 1-2 and 13-14). Thus, MCM10 promotes total DNA replication, consistent with (Wohlschlegel et al. 2002), but is not required for DNA replication in vertebrates, consistent with (Chadha et al. 2016).

**Supplemental Figure 3:**
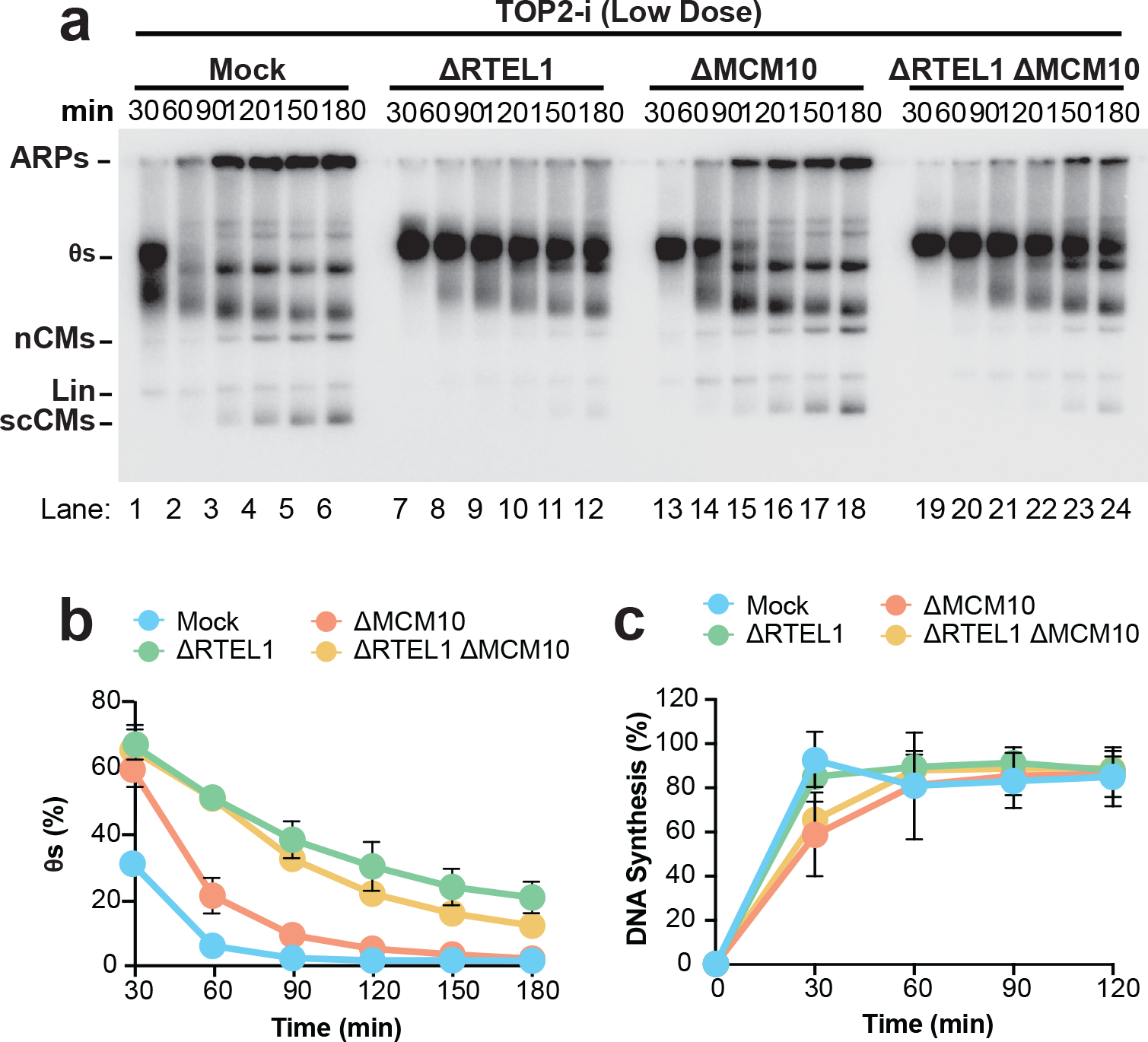
Co-depletion of RTEL1 and MCM10 does not impact total DNA replication. (A) Replication was performed as in Figure 3A and replication intermediates were analyzed directly, without restriction digest. Catenanes (Cats) (Dewar et al. 2015) and aberrant replication products (ARPs) (Deng et al. 2018; Heintzman et al. 2019) were previously defined. (B) Quantification of θ structures from (A) as a read out for fork merger. (C) Total DNA synthesis from (A) was measured. Mean ± SD, n=3 independent experiments

**Supplemental Figure 4:**
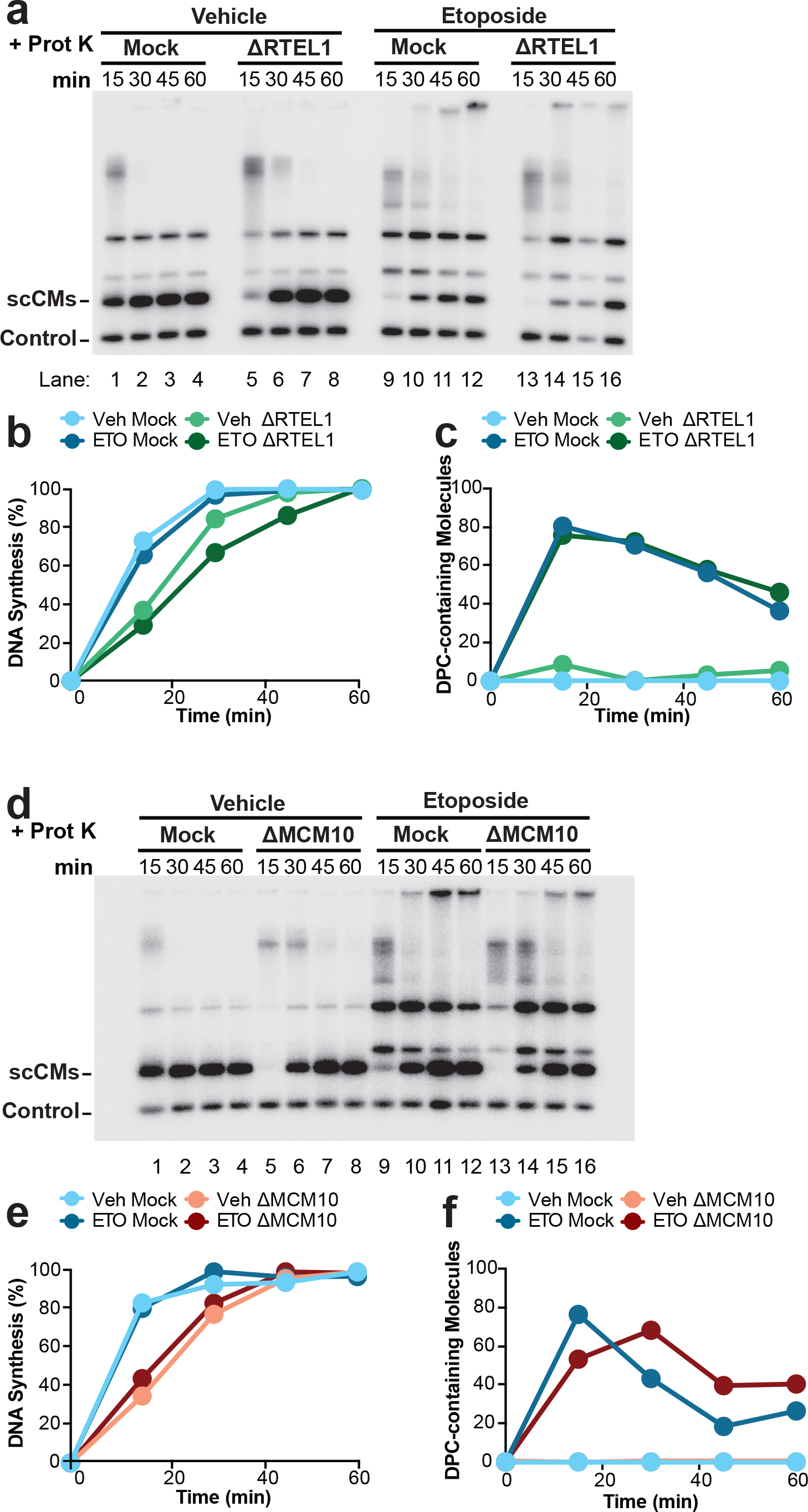
Analysis of replication intermediates formed following etoposide treatment. (A) Autoradiogram of proteinase K-treated samples corresponding to the untreated samples in Figure 4B. (B) Total DNA synthesis was measured from (A), normalized to the control. (C) An experimental replicate of Figure 4C (D) Autoradiogram of proteinase K-treated samples corresponding to the untreated samples in Figure 4D. (E) Total DNA synthesis was measured from (D), normalized to the control. (F) An experimental replicate of Figure 4E

**Supplemental Figure 5:**
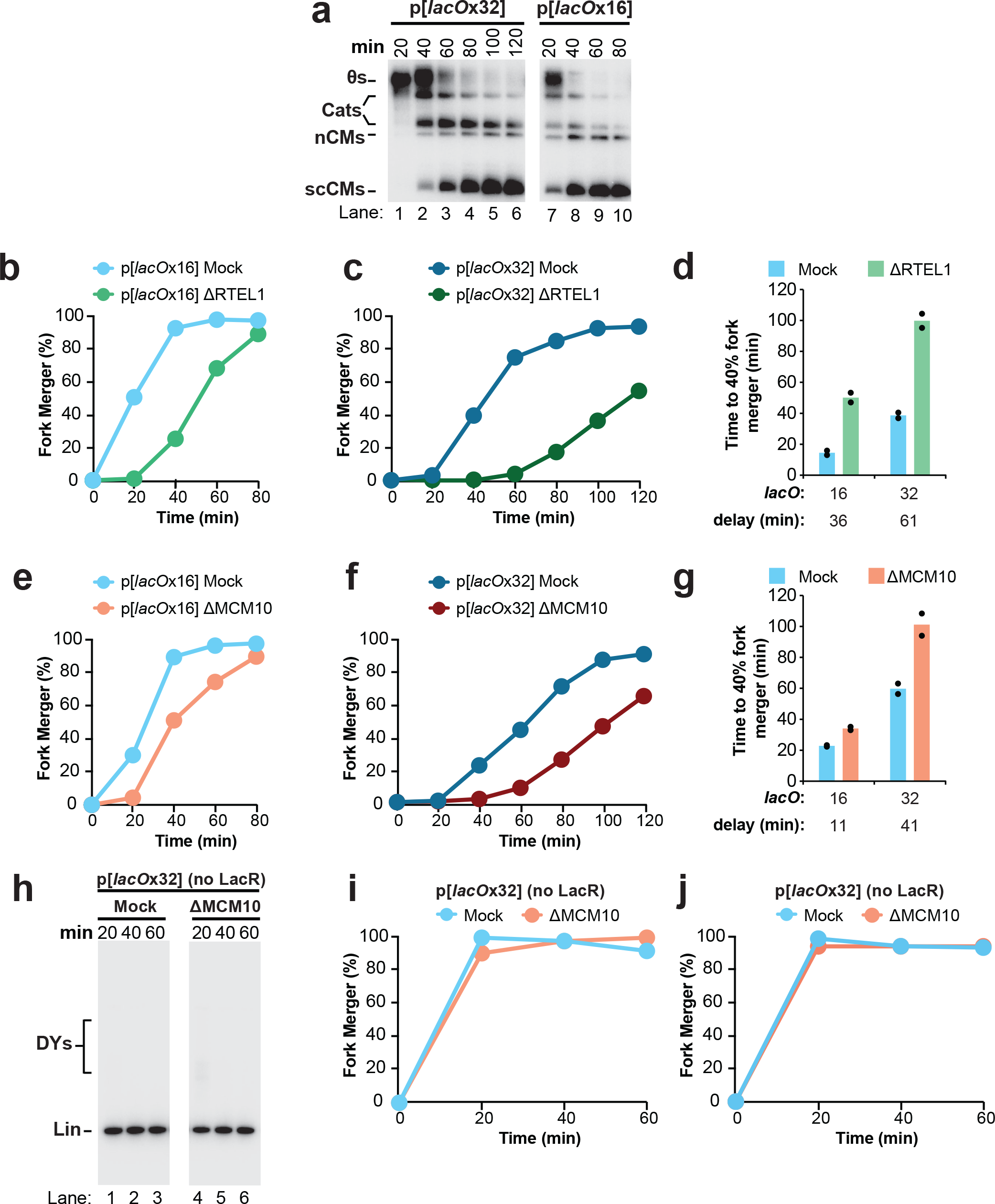
Analysis of fork progression through a LacR barrier. (A) Replication was performed as in Figure 5A using mock-depleted extracts and replication intermediates were analyzed directly, without restriction digest. Catenanes (Cats) were previously defined in (Dewar et al. 2015). (B) An Experimental replicate of Figure 5C (C) An Experimental replicate of Figure 5E, which is part of the same experiment shown in (A) (D) The time at which 40% of forks merged was calculated for Figure 5C,E and (B)-(C). Means were plotted (bars) alongside the individual values (black circles). Mean values were used to calculate the delay in fork merger caused by RTEL1 depletion (ΔRTEL1) compared to the control (mock). (E) An Experimental replicate of Figure 5G (F) An Experimental replicate of Figure 5I, which is part of the same experiment shown in (D) (G) The time at which 40% of forks merged was calculated for Figure 5G,I and (E)-(F) as in (D). Mean values were used to calculate the delay in fork merger caused by MCM10 depletion (ΔMCM10) compared to the control (mock). (H) As part of the experiment shown in Figure 5F,H fork merger was monitored in mock and MCM10-depleted extracts as in Figure 5A but in the absence of LacR. Samples were separated on a native agarose gel and visualized by autoradiography. (I) Fork merger was measured from (H). (J) An experimental replicate of (I).

